# A novel machine-learning classification model detects oxidative fiber type transitions in a rabbit model of cerebral palsy

**DOI:** 10.64898/2026.06.11.731759

**Authors:** Cassandra Kramer, Emily J. Reedich, Hope McCann, Sadie Drouin, Destiny Sanders, Elian Gonzalez, Tiffany Ung, Alexandra Mukisa, Elvia Mena Avila, Brendan Moline, Landon Genry, Jess Glennon, Camila Quiroga, Lisa Dowaliby, Christine J. DiDonato, Katharina A. Quinlan, Marin Manuel

## Abstract

The distribution of slow-and fast-twitch fiber types in a skeletal muscle heavily influences its physiology. Muscle biopsy studies indicate atypical fiber type composition and fiber size variation in children with cerebral palsy (CP), but subjects have variable treatment history and a variety of muscles affected, so uncertainties remain. In this study, we developed a novel machine-learning classification model to perform high-throughput fiber typing of complete transverse muscle sections. Our XGBoost algorithm-based prediction model yielded a balanced accuracy score of 0.89 and a macro F1-score of 0.89, reflecting its ability to robustly predict muscle fiber type from myosin heavy chain (MyHC) isoform immunofluorescence intensities and morphological descriptors. This is the first reported fiber type classifier to consider hybrid fibers, which is a major advance, considering at least 20% of myofibers are hybrid yet they are routinely overlooked due to difficulty in their detection. We used this classification model to define fiber types of more than 7 million myofibers from flexor-extensor muscle pairs in rabbits that experienced hypoxia-ischemia (HI) injury in utero (modeling CP), and typically developing sham rabbits. We observed an oxidative fiber type shift in flexor muscles (biceps brachii and tibialis anterior) of HI rabbits at postnatal day (P)14-20 and P30-32 (weaning age). This altered fiber type composition imparts reduced contractile force and is amenable to sustained muscle activity; it may reflect chronic low-frequency motor unit activation. This work supports prior clinical reports that developmental trajectories of muscle fibers are disrupted in CP.

## Background

The developmental disorder cerebral palsy (CP) is the most common cause of lifelong motor impairment. CP-causative brain injuries are perinatal and occur during a critical period for neuromuscular system development. Reflexive and volitional movements are dysfunctional in CP (Capute 1979; Milner-Brown and Penn 1979; Poon and Hui-Chan 2009; Sukal-Moulton and Fowler 2020), and the signaling relays that underlie these motor processes conclude with excitation-contraction coupling in skeletal muscle. Muscle fibers are classified into physiological types based on shortening velocity, force-generative capacity, and metabolism. These properties exist on a spectrum and are dictated by the isoform of myosin heavy chain (MyHC) expressed in sarcomeric thick filaments and the characteristic Ca^2+^-ATPase activity it confers for cross-bridge cycling (Pette and Staron 2000; Schiaffino et al. 2025; Schiaffino and Reggiani 2011). Fiber type composition imparts certain physiological characteristics to every muscle, with important functional consequences. Type I muscle fibers are encoded by the MYH7 gene; these myofibers slowly contract (they are labeled ‘slow-twitch’), generate relatively weak forces, and are prominent in postural muscles. Since the primary metabolic pathway used by type I myofibers is oxidative phosphorylation, these fibers are resistant to fatigue. Type IIa myofibers express MYH2; these fast-twitch fibers generate moderate forces and are largely fatigue-resistant. Type IIx muscle fibers are encoded by MYH1; they are fast-twitch, generate relatively large forces, and are susceptible to fatigue because of primarily glycolytic metabolism, although they are much more oxidative in rodents than in humans (Schiaffino 2010). Type IIb muscle fibers are encoded by MYH4, which turns on last in development, and occupy the far-end of the spectrum. These glycolytic fast-twitch fibers contract the fastest, generate the strongest forces, and are highly fatigable. MYH4 is not typically expressed in human muscle. Muscle fiber type is plastic. Hybrid fibers express more than one MyHC isoform, and various conditions like exercise, disuse, disease, and injury can modulate MyHC expression in muscle cells (Medler 2019).

Children with CP exhibit altered muscle fibers. For instance, several studies have shown an increase in muscle fiber size variation in type I and/or type II fibers (Deschrevel et al. 2024; Deschrevel et al. 2026; Ito et al. 1996; Kahn et al. 2023; Rose et al. 1994), which has been associated with increased electromyographic activity and greater energy expenditure during walking (Rose et al. 1994). Increased fiber size variation has been observed in other neuromuscular disorders and is not a pathology specific to spastic muscle (Dubowitz and Brooke 1973; Lieber et al. 2004; Valentine 2017). Muscle fiber type compositions are also altered in children with CP, but the pattern in which they are changed is not well characterized. One factor at play may be that muscle biopsies do not necessarily represent muscles at-large. A second consideration is that the experimental design of histological or biochemical studies has sometimes involved ill-fitting control groups or pooled data from disparate muscles (reviewed in (Edman et al. 2026)), presumably because it is inherently challenging to obtain consent for muscle biopsies from young children.

In a well-controlled histological study, Robinson et al. (2013) examined fiber type composition of erector spinae biopsies obtained from scoliotic children with and without spastic CP, all who were undergoing spinal fusion surgery; they reported an increased proportion of type I and decreased proportion of type II fibers in the CP condition (Robinson et al. 2013). Most recently, Deschrevel et al. observed increased proportions of type IIx fibers in gastrocnemius and semitendinosus muscle biopsies (paired with a decreased type I proportion in the former) from school-aged children with spastic CP but saw minimal or no differences in samples from even younger children (Deschrevel et al. 2024; Deschrevel et al. 2026). However, confounding factors that add variability in these studies are disuse and muscle tone-treatment history. Disuse minimally affects predominantly fast-twitch muscles but causes slow-to-fast fiber type transition in mostly slow-twitch muscles (Gupta et al. 1989; Talmadge et al. 1996); botulinum neurotoxin type-A (Botox) injections, a treatment for spasticity, is known to cause slow-to-fast fiber type transitions (Korfage et al. 2012; Valentine et al. 2016). Additional molecular biological analyses have yielded evidence for slow-to-fast fiber type transitions in wrist flexors and extensors (Gantelius et al. 2012; Smith et al. 2009) and slower profiles in the gracilis and semitendinosus muscles in CP (Smith et al. 2012; Smith et al. 2011). Taken together, these findings suggest that muscle fiber type transitions in CP are likely age-dependent and muscle-specific, although interpretation is complicated by confounding effects of disuse, treatment history, and limitations inherent to muscle biopsies.

One risk factor for CP is perinatal hypoxia-ischemia (HI) injury (Graham et al. 2008; Robertson and Finer 1985), which occurs in 6 of 1000 live births (Madan et al. 2005).The goal of our study was to measure muscle fiber type composition in skeletal muscle of young rabbits that experienced HI injury in utero. We chose to conduct our study using the HI rabbit model of CP because rabbits show several clinically relevant CP-like features after developmental injury, which is unlike the rodent (reviewed in (Cavarsan et al. 2019; Quinlan et al. 2026). To achieve this goal, we developed a machine-learning prediction model for high-throughput classification of muscle fiber type from MyHC immunofluorescence intensity and cell morphometry. We used our XGBoost algorithm-based classification model to define fiber type transition patterns in flexor-extensor muscle pairs of HI rabbits at two stages of postnatal development, postnatal day (P)14-20 and P30-32 (weaning age). We found that at both ages, prenatal HI injury imparted a more oxidative fiber type profile to the biceps brachii and tibialis anterior (flexor muscles). This fiber type transition may reflect immaturity at P14-20 and chronic low-frequency motor unit activity at P30-32. Functionally, it would facilitate sustained muscle activity and hypertonia.

## Methods

### Animal housing and husbandry

New Zealand white rabbit kits used in this study were born in a controlled vivarium with a 12:12 h light-dark photoperiod to dams that had access to food and water ad libitum. Dams were either bred in-house or obtained as timed-pregnant from Charles River Laboratories, Inc. (Wilmington, MA). All rabbits in our breeding program originated from Charles River Laboratories.

### Surgical procedures

Surgery was performed on pregnant dams to induce global HI in fetal rabbits while in utero. The surgical procedure was performed during late gestation as described previously (70-92% gestation; embryonic day (E)22-29 of 31.5 days gestation; (Derrick et al. 2004; Reedich et al. 2023; Reedich et al. 2022; Steele et al. 2020)). Rabbits were pre-medicated with ketamine (25 mg/kg) and xylazine (1.0-3.75 mg/kg) or “bunny magic” (5 mg/kg ketamine; 0.05 mg/kg buprenorphine; 0.025 mg/kg dexdomitor) administered by intramuscular injection. Rabbits were anesthetized with isoflurane (1-5%), heart rate was measured by electrocardiogram, oxygen saturation and respiration were measured by pulse oximeter. Body temperature was monitored using a rectal thermometer and maintained between 101 °F and 103 °F using heat support. An adequate surgical plane was confirmed by a stable heart rate of 150-250 beats per minute (BPM) and the absence of withdrawal reflex to toe pinch. Rabbits that showed elevated heart rate (over 250 BPM or increased respiratory rate during the procedure received additional doses of buprenorphine or ketamine. The femoral artery was isolated and a Fogarty balloon catheter (2-3 French) (Edwards Lifesciences, cat. # 120602F, 120403F) was inserted and advanced into the femoral artery so that the balloon was positioned just rostral to the bifurcation of the descending aorta. The balloon was inflated with saline for 40-45 minutes to occlude circulation to the uterine horns. Placement of the catheter and vessel occlusion was confirmed by ultrasound. Sham surgical procedures followed the same protocol but without insertion or inflation of the catheter. Dams recovered after surgery and later gave birth to kits at full-term on the expected due date ± 2 days. Forty-eight rabbit kits (24 sham and 24 HI; mixed sex) were used in this study. We grouped together male and female kits because we evaluated muscle fiber type composition in kits much younger than the age at which they reach sexual maturity (∼6 months of age).

### Motor behavioral testing

Newborn kits are not equally affected by the HI surgical procedure that they experienced in utero (Derrick et al. 2004; Shi et al. 2022). Therefore, every kit is tested at birth. Ashworth testing: We classified newborn HI kits by the maximum hypertonia score that they displayed in the early postnatal period, as previously described (Derrick et al. 2004; Reedich et al. 2022). Briefly, we used a modified Ashworth score developed for rabbits to quantify the degree of hypertonia displayed by HI kits at time points from postnatal day (P) 1-18. Hypertonic HI kits had muscle tone scores of ≥3 in one or more evaluated joints; muscle tone scores were 1–2 in all joints of non-hypertonic HI kits, indicating a normal amount of voluntary resistance with no spastic catch. Open field testing: At P1, rabbit kits were placed into a clear plexiglass box (45 cm x 30 cm) with a black rubber mat at the bottom. Rabbit kits were allowed to explore the box for two minutes. During testing, videos were recorded using a 1080p webcam and data was collected with ANY-maze software (version 7.65, Stoelting Co.). The immobility detection threshold was set to 65% and the minimum immobility period was set to 2000 ms. The immobility detection threshold differentiates between locomotion, moving from one place to another, and smaller movements such as grooming. Non-hypertonic and hypertonic HI kits were pooled for subsequent analysis and hypertonia as a biological variable was evaluated post-hoc.

### Tissue processing

Sham and HI rabbits were euthanized in the third postnatal week (P14-20), hereafter called the “P14 age group”, or at weaning-age (P30-32), hereafter called the “P31 age group”, by overdose of sodium pentobarbital; decapitation as a secondary measure of euthanasia was implemented. As the dissection progressed, fur over the fore-or hind-limb was wetted with alcohol and the fur was removed. The following flexor-extensor muscle pairs were dissected and collected onto phosphate buffered saline (PBS)-wetted gauze: the tibialis anterior, soleus, gastrocnemius (medial and lateral heads), plantaris, biceps brachii, and triceps brachii (medial, long, and lateral heads). All muscles were fresh-frozen in liquid nitrogen-chilled isopentane (∼-160 °C) and stored at-80 °C until cryosectioning. Transverse sections from each muscle mid-belly were collected at 12-µm thickness onto 8 microscope slides such that for any given slide, the distance between adjacent slices was ∼96 µm. Slices placed consecutively on adjacent slides were therefore only ∼12 µm apart and sampled the same population of myofibers. Slides were stored at-80 °C until immunofluorescence.

### Immunofluorescence

For each muscle, one pair of adjacent slides were selected for fluorescent labeling of extracellular matrix by fluorophore-conjugated wheat-germ agglutinin (WGA) plus immunolabeling of MyHC isoforms expressed in myofibers. This staining protocol is adapted from Bloemberg and Quadrilatero (2012) (Bloemberg and Quadrilatero 2012). Selected slides were thawed, briefly rehydrated in PBS, and blocked for 1 h in blocking buffer (composition: 5% goat serum (Jackson ImmunoResearch, cat. # 005-000-121), 0.5% Triton X-100 (Sigma-Aldrich, cat. # X100-5ML), and 0.5% glycine (Sigma-Aldrich, cat. #G7403-100G) in PBS). After blocking, incubations were performed overnight at 4 °C with mouse primary antibody cocktails diluted in blocking buffer. The first cocktail was against MyHC-I (1:50, DSHB, cat. # BA-F8; isotype IgG2b), MyHC-IIa (1:00, DSHB, cat. # SC-71; isotype IgG1), and MyHC-IIb (1:100, DSHB, cat. # BF-F3; isotype IgM). Slices on the subsequent slide were incubated with the second primary antibody cocktail against MyHC-IIa (1:200, DSHB, cat. # SC-71; IgG1) and MyHC-IIx (1:20, DSHB, cat. # 6H1; isotype IgM). Afterwards, both slides were washed in 0.1% Tween-20 (Sigma-Aldrich, cat. # P9416-100ML) in PBS (PBST). Then, incubations were performed for 2 h at room temperature with isotype-specific secondary antibody cocktails diluted in blocking buffer. The first cocktail contained DyLight™-405 Goat anti-Mouse IgG2b (1:500, Jackson, cat. # 115-475-207), Alexa Fluor™-488 Goat anti-Mouse IgG1 (1:500, Life Technologies, cat. # A21121), and Alexa Fluor™-594 Goat anti-Mouse IgM (1:500, Jackson, cat. # 115-585-075). The subsequent slide was incubated with the second cocktail containing Alexa Fluor™-488 Goat anti-Mouse IgG1 (1:500, Life Technologies, cat. # A21121) and Alexa Fluor™-594 Goat anti-Mouse IgM (1:500, Jackson, cat. # 115-585-075). After washing with PBST, both slides were incubated for 2 h at room temperature with WGA-Alexa Fluor™-647 (1:100 diluted in PBS, ThermoFisher, cat. # W32466) to label extracellular matrices. After washing with PBST, both slides were rinsed once quickly in PBS and mounted with number 1.5 coverlips (VWR # 48393-241) using Fluoromount (Sigma-Aldrich, cat. # F4680-25ML). Once dry, slides were sealed with nail polish and stored at-20 °C until microscopy. For every staining cohort, one negative control slide was included to confirm the absence of any non-specific fluorescent labeling from the triple-secondary antibody cocktail; the negative control received only blocking buffer in place of primary antibody cocktail and was treated the same as the experimental slides for all other steps.

### Epifluorescent microscopy

Widefield epifluorescent photomicrographs were acquired on a Nikon ECLIPSE Ti2-E inverted microscope equipped with a motorized stage and a Sola light engine (Lumencor). Observations were performed using a Nikon CFI Plan Apochromat Lambda D 10×/0.45NA air objective, a DAPI Filter Set (Nikon #96360, excitation: 350/50, dichroic: 400, emission: 460/50) for DyLight™ 405, a GFP Filter Set (Nikon #96362, excitation: 470/40, dichroic: 495, emission: 525/50) for Alexa Fluor™ 488, a DsRed/TRITC/Cy3 Filter Set (Nikon #96394, excitation: 554/23, dichroic: 573, emission: 609/54) for Alexa Fluor™ 594, and a Cy5 filter set (Nikon #96232, excitation:618/50, dichroic: 652, emission: 698/70) for Alexa Fluor™ 647. Images were acquired using a Zyla 4.2 sCMOS camera (Andor; 12-bit, gain 4, rolling shutter, binning 2×2) and NIS-Elements AR v.6.10.01 (Nikon). Light intensity and exposure time were adjusted to maximize the dynamic range of each image.

Before analysis, each image was processed in a standardized way, using ImageJ (v. 2.16.0/1.54p). First, we applied a rolling ball background subtraction (ball diameter 200px) on each channel; then, if the sample contained several muscles or muscle heads, each muscle was cropped and saved as a separate file; finally, minor blemishes (small folds, bubbles) were deleted manually from the files. The WGA channel was separated and saved separately, while the 3 channels of fluorescence from MyHC-I, MyHC-IIa, and MyHC-IIb were saved together in a 16bit TIFF file for subsequent analysis.

### Prediction model

Each WGA image file was segmented using cellpose-SAM (v.4.1.1 (Pachitariu et al. 2025)). The masks were then applied to the 3-channel fluorescence image and the following measurements were taken using scikit-image (v0.26.0 (Walt et al. 2014)): cross-sectional area (CSA), fiber perimeter, eccentricity (eccentricity of the ellipse that has the same second moments as the region, in the range [0-1], where 1 is a circle), equivalent diameter area (the diameter of a circle with the same area as the region), major and minor axis lengths (lengths of the major and minor axis of the ellipse that has the same normalized second central moments as the region), axis ratio (ratio of the major and minor axis lengths), roundness (calculated as r = 4π x area / perimeter^2^, in the range [0-1] where 1 is a perfect circle), maximum Feret’s diameter (computed as the longest distance between points around a region’s convex hull), and the mean intensity of the fluorescence in each of the 3 channels.

For model training, we generated a training set by randomly selecting 2116 muscle fibers sampled from P14 and P31 rabbit gastrocnemius for manual fiber type classification. Images that depicted the MyHC-I, MyHC-IIa, and MyHC-IIb immunofluorescence of these myofibers (leaving MyHC-IIx unstained) were used to visually determine muscle fiber type. Images that showed MyHC-IIa and MyHC-IIx immunofluorescence on the subsequent cryosection, sampled 12-µm away, were used to check these same cells for MyHC-IIx signal. This was done to confirm that low intensity DyLight™ 405, Alexa Fluor™ 488, and Alexa Fluor™ 594 signals - or the absence of any staining - in these fibers was properly interpreted as hybrid or pure MyHC-IIx expression, respectively. We noted that the mouse anti-MyHC-IIx primary antibody (DSHB, cat. # 6H1) strongly labeled type IIx fibers on the subsequent slide but also bound to type I fibers with low affinity, an artifact that has also been observed by others (Rehman et al. 2025).

The final model was implemented in scikit-learn (v.1.8.0 (Pedregosa et al. 2011)) using XGBoost (version 3.2.0) with a multi-class classification objective (multi:softprob) to predict six muscle fiber classes. Hyperparameters were optimized via randomized search with cross-validation. The selected model used max_depth = 4, learning_rate = 0.05, n_estimators = 300, min_child_weight = 10, gamma = 2.0, subsample = 0.6, colsample_bytree = 0.8, reg_alpha = 0.5, reg_lambda = 1.0. The model was trained using the histogram-based tree construction algorithm (tree_method = hist).

## Statistical analysis

All statistical analyses were performed using Python (version 3.14.5) with standard scientific libraries (Numpy v.2.4.6, Pandas v.3.0.3, Statsmodel v.0.14.6, matplotlib v.3.10.9, seaborn v.0.13.2).

### Open field

P1 kit behavior during the open field test was measured by the total distance traveled, the total time spent mobile, and the average speed. We tested whether these variables were different between shams, HI kits that were categorized as hypertonic based on Ashworth score, and HI kits that were not hypertonic. For average speed and total distance traveled, we performed a one-way ANOVA after checking residual normality and homogeneity of variance. For total time mobile, these assumptions were not met, so group differences were assessed using Welch’s ANOVA. To account for multiple tests, p-values were adjusted using the Benjamini–Hochberg false discovery rate procedure. Post-hoc pairwise comparisons were performed using Tukey’s HSD test for variables analyzed by standard ANOVA and Games-Howell tests for the variable analyzed by Welch’s ANOVA.

### Image analysis, data inclusion and preprocessing

Manual quality assessment indicated a segmentation error rate of 3.4% by cellulose-SAM (see Results), primarily due to merged fibers, inclusion of non-fiber regions, or segmentation artifacts. To minimize the influence of these errors while avoiding manual bias, we excluded fibers with cross-sectional area in the upper and lower 2% of the distribution, resulting in removal of 4% of fibers per sample. This threshold was chosen because it closely matches the empirically measured segmentation error rate (3.4%). All remaining fibers were included in downstream analyses without additional manual curation (see Table 1 for total counts).

### Fiber type composition analysis

Differences in fiber type proportions between groups (HI vs. sham), ages (P14 vs. P31 age groups), and muscles were assessed using linear regression models applied to animal-level summary data. For each animal and muscle, proportions of each fiber type were computed and used as response variables.

To estimate group differences, we fit linear models of the form proportion ∼ group for each age group, muscle and fiber type. Estimated differences between HI and sham groups (Δ = HI − sham) were extracted from model coefficients. Because of potential heteroskedasticity, robust standard errors (HC3) were used for calculations. For each comparison, 95% confidence intervals were computed. To account for multiple testing across fiber types, muscles, and age groups, p-values were adjusted using the false discovery rate (FDR) procedure (Benjamini–Hochberg). Adjusted p-values are reported as q-values.

### Fiber size and variability analysis

Muscle fiber cross-sectional area and coefficient of variation (CV) were analyzed using linear mixed-effects models to account for the hierarchical structure of the data (fibers nested within animals and muscles). Models included fixed effects for group (HI vs. sham), age groups (P14 vs. P31), fiber type, and their interactions, with random intercepts for animal identity. The general model structure was: Y ∼ group * age * fiber type + (1 | animal) where *Y* is the target variable (area, or CV of area) and (1 | *animal*) is a random intercept per animal.

Model assumptions were checked by visual inspection of residuals (QQ plot). Statistical significance of fixed effects was evaluated using Wald tests, and results are reported with estimated effect sizes, 95% confidence intervals, and p-values.

## Results

### A statistical prediction model for high-throughput fiber type determination

We set out to establish a method for unbiased classification of muscle fibers across entire muscles. To achieve this, we developed a pipeline combining automated image segmentation with supervised machine learning (Figure 1A).

**Figure 1:**
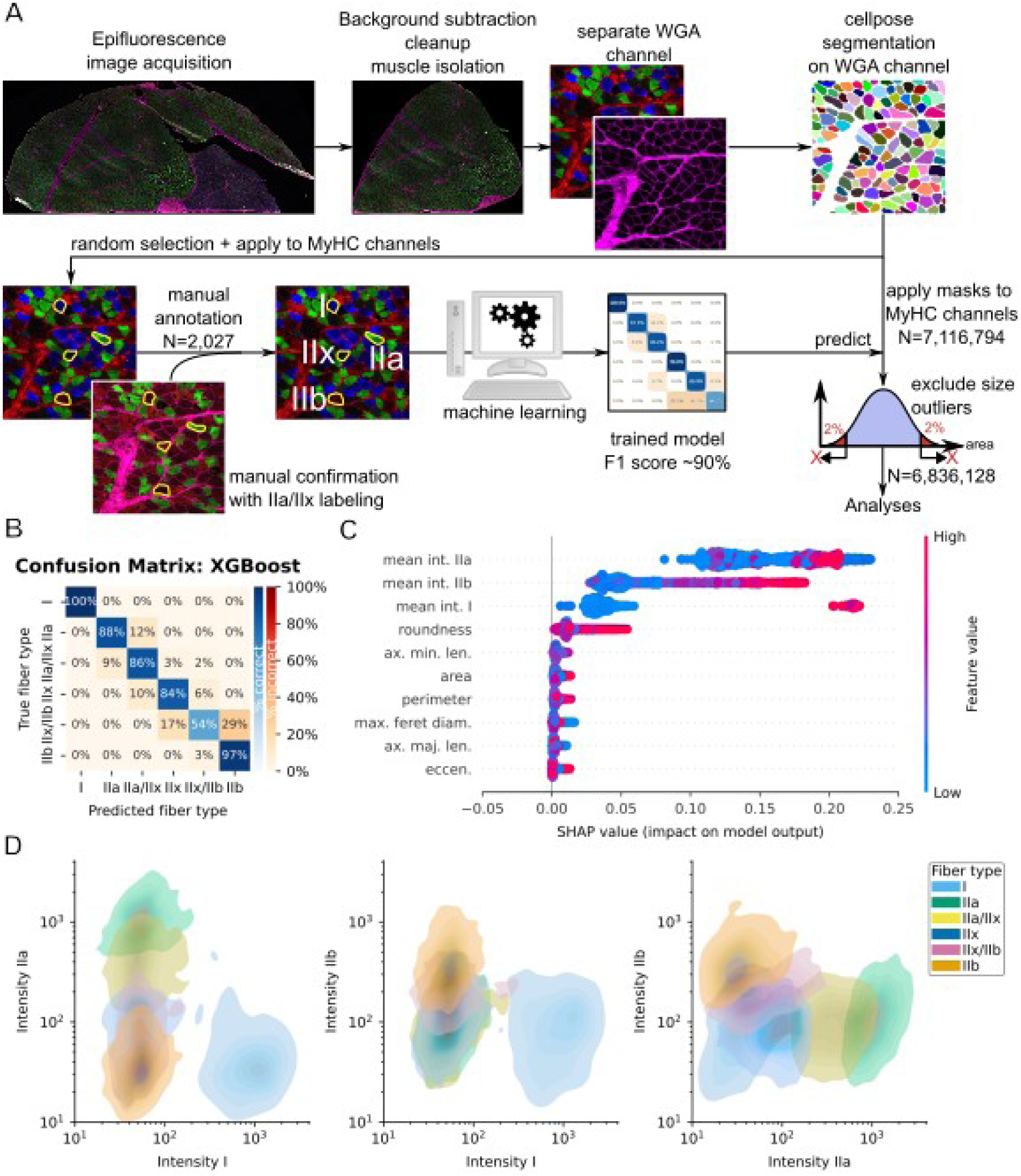
Fiber type prediction model. **(A)** Schematic representation of the study workflow. **(B)** Normalized confusion matrix for the six classes (I, IIa, IIa/IIx, IIx, IIb, IIx/IIb). The main diagonal (blue) shows the proportion of fiber types correctly classified, while the off-diagonal squares (red-orange) show the proportion of units that were misclassified. **(C)** Shapley Additive Explanations (SHAP) summary plot showing the global importance and effect of input features on model predictions. Each point represents an individual muscle fiber, with position on the x-axis indicating the magnitude and direction of the feature’s contribution to the model output (SHAP value). Features are ranked by overall importance from top to bottom. Color represents the feature value (blue = low, red = high). Abbreviations: mean int. IIa: average intensity of MyHC-IIa signal. mean int. IIb: average intensity of MyHC-IIb signal. mean int. I: average intensity of MyHC-I signal. ax. min. len.: minor axis length. max. ferret diam.: maximum Feret’s diameter. ax. maj. len.: major axis length. eccen.: eccentricity. **(D)** Distribution of the fluorescence intensity in each of the three channels (MyHC-I, MyHC-IIa, MyHC-IIb channels) in the final dataset. Each fiber type is represented by a series of color-coded contours obtained by bivariate kernel-density estimation. The graphs demonstrate clear separation between fiber types. Type I fibers cluster at high MyHC-I signal, type IIb fibers at high MyHC-IIb signal, and type IIa fibers occupy intermediate regions. Axes are shown on a logarithmic scale of 12-bit fluorescence values.

First, all muscle images were segmented using cellpose-SAM (Pachitariu et al. 2025), applied to the WGA fluorescence channel that delineates muscle fiber boundaries. The masks were then applied to the immunofluorescence channels of the MyHC immunolabeling and the morphological measurements (e.g., area, circumference, circularity), as well as the average fluorescence intensity for each of the three channels (MyHC-I,-IIa,-IIb) was measured inside each mask.

From the total dataset, approximately 2100 muscle fibers were randomly selected across multiple muscles to generate a labeled dataset. Each fiber was independently reviewed by two expert investigators, who manually identified muscle fiber type based on immunofluorescence intensity of MyHC isoforms (MyHC-I,-IIa, and-IIb), as well as MyHC-IIx expression assessed from a serial section located 12 µm apart. Manual quality assessment indicated that cellpose correctly segmented 96.6% (N=2044) of the muscle fibers. Only 3.4% of segmented objects were incorrect, primarily due to inclusion of connective tissue regions, merging of adjacent fibers, or other segmentation artifacts. This low error rate demonstrated that cellpose provided highly accurate segmentation, enabling inclusion of essentially all fibers within each muscle and thereby avoiding the sampling bias inherent to manual selection approaches.

As expected based on the natural distribution of muscle fibers in muscles, the distribution of fiber types in the annotated dataset was imbalanced, with the majority of fibers corresponding to type IIa (N=651), type IIb (N=414), and hybrid type IIa/IIx fibers (N=386), followed by type IIx (N=210), type I (N=182), and hybrid type IIx/IIb (N=160). Rare hybrid classes, including I/IIa (N=12), IIa/IIb (N=4), and IIa/IIx/IIb (N=8), were excluded from model training due to insufficient sample size for reliable learning.

From this dataset, 300 fibers were held out as an independent test set, while the remaining ∼1700 fibers were used for model training and hyperparameter tuning. Multiple supervised learning algorithms were evaluated, with hyperparameters optimized via grid search using cross-validation. Because fiber type proportions are imbalanced, model performance was assessed using balanced accuracy.

We evaluated multiple supervised learning algorithms for fiber type classification, including linear models (logistic regression, linear discriminant analysis), support vector machines, and tree-based ensemble methods (random forest, extra trees, gradient boosting, and XGBoost). Model performance was assessed using balanced accuracy and macro F1-score on the held-out test set, due to class imbalance across fiber types. Among conventional models, ExtraTrees achieved the strongest performance, with a test accuracy of 0.863, balanced accuracy of 0.853, and macro F1-score of 0.847. Random forest and histogram-based gradient boosting also performed well, with balanced accuracies of 0.822 and 0.831, respectively. XGBoost models consistently outperformed all other approaches. The best-performing model achieved an accuracy of 0.90, balanced accuracy of 0.89, and macro F1-score of 0.89 on the held-out test set (N=300 fibers). XGBoost was selected because it provided the highest balanced accuracy and macro F1-score, indicating superior performance across all fiber classes, including minority classes.

The final XGBoost model showed strong performance across most fiber types (Figure 1B). Pure fiber types were classified with high precision and recall, particularly type I fibers, which were perfectly classified (precision = 1.00, recall = 1.00), and type IIb fibers (precision = 0.90, recall = 0.97). Type IIa fibers also showed high performance, although with slightly lower recall (0.88). Not surprisingly, the model showed reduced performance for hybrid fiber types, with type IIa/IIx and type IIx/IIb exhibiting lower precision (0.77 and 0.72, respectively) and recall (0.86 and 0.54, respectively). In particular, the type IIx/IIb class showed the greatest difficulty, reflecting substantial overlap with neighboring fiber types. Nevertheless, even when misclassifying fibers, the model generally ascribed them to a related fiber type. For instance, misclassified type IIa/IIx fibers might be labeled as type IIa or type IIx, but never type I; and the type IIx/IIb fibers that are misclassified ended up labeled as either type IIx or IIb, but not type IIa or I.

To understand the role of the features in the model, we used Shapley Additive Explanations (SHAP) (Lundberg and Lee 2017), which quantifies the contribution of each feature to model predictions. This approach enabled us to compare the relative importance of immunofluorescence intensity measurements and morphological descriptors in determining muscle fiber type. The SHAP analysis (Figure 1C) demonstrates that the model’s decision-making is dominated by fluorescence intensity; the relationships between MyHC fluorescence intensities and fiber type are depicted in Figure 1D. Morphological features contribute only marginally; the morphological descriptor with the highest impact was roundness. This confirms that the classifier faithfully uses established criteria for muscle fiber typing rather than relying on indirect or confounding features.

### Flexor muscles are more oxidative after prenatal HI injury

The main purpose of this study was to define how prenatal HI injury (a risk factor for CP) disrupts the establishment of muscle fiber type composition in early postnatal development. We used our novel fiber type classification model to quantify fiber type compositions of flexor-extensor muscle pairs acting at the ankle and elbow, joints that are sometimes hypertonic in neonatal rabbits that experienced HI injury in utero (Figure 2). We first measured fiber type compositions in the third postnatal week (“P14 age group”), which represents a dynamic developmental stage. At this age, synapse elimination at poly-innervated neuromuscular junctions is complete (Bixby and Van Essen 1979), kits bear weight, and have begun displaying mature locomotor patterns like hopping.

**Figure 2:**
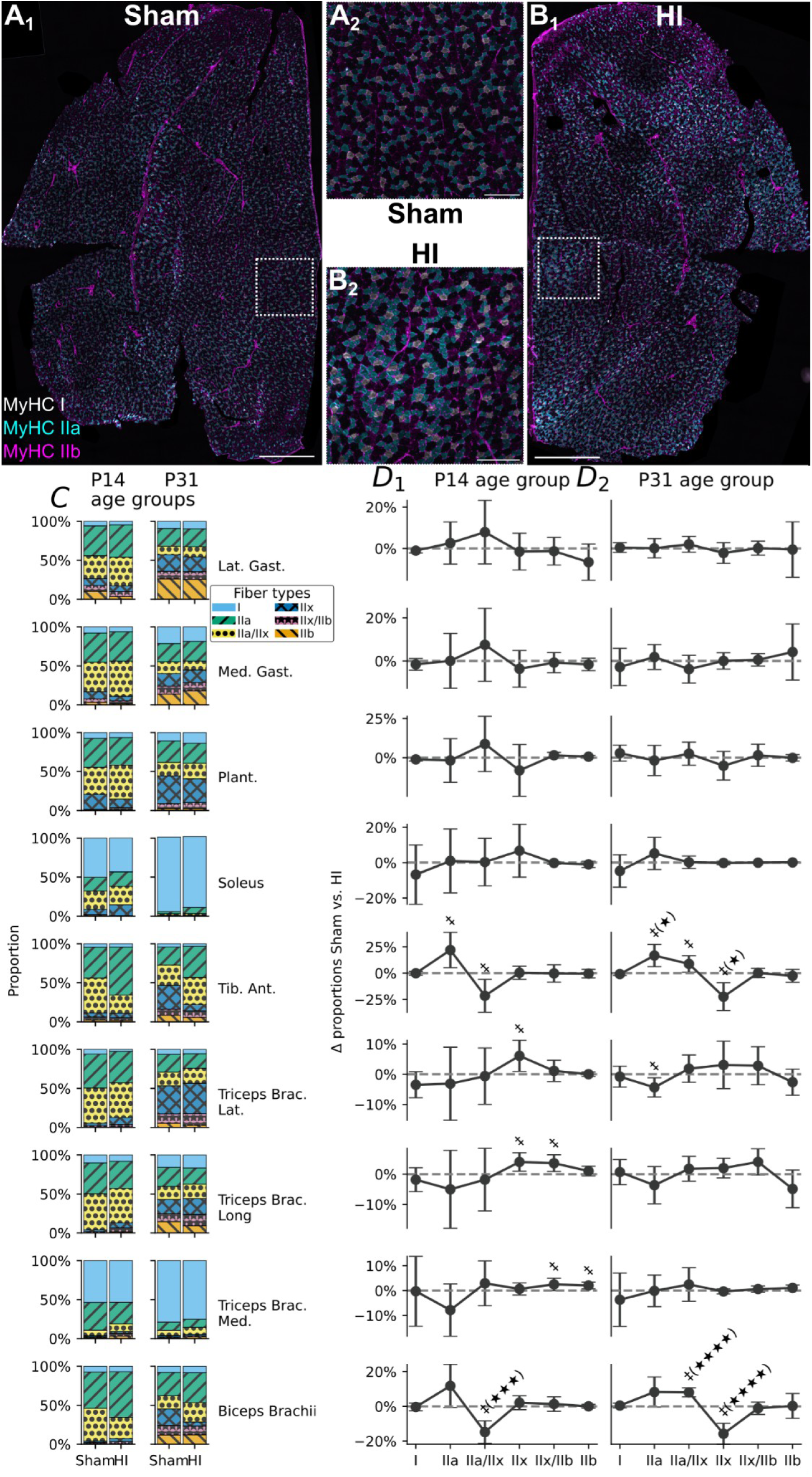
Fiber type composition. (A_1_) Representative image of a cross section of a biceps muscle from a P31 sham rabbit, stained for MyHC-I (white), MyHC-IIa (cyan) and MyHC-IIb (magenta). Scale bar: 1 mm. **(A_2_)** Zoom on the region highlighted in A_1_. Scale bar: 200 µm. **(B_1_)** Representative image of a cross section of a biceps muscle from a P31 HI rabbit. Same stain as in A_1_. Scale bar: 1 mm. **(B_2_)** Zoom on the region highlighted in B_1_. Scale bar: 200 µm. **(C)** Stacked histograms showing the distribution of muscle fibers I, IIa, IIa/IIx, IIx, IIx/IIb, and IIb in each studied muscle as a function of group (sham vs. HI) and age (P14 vs. P31 age group). **(D)** Estimated differences in fiber type proportions between HI and sham groups (Δ proportion; HI − sham) are shown across muscles and age groups. Panels are organized by muscle (rows) and age (columns: P14 age group **(D_1_)** and P31 age group**(D_2_)**). Points represent model-based estimates from linear regression, and error bars indicate 95% confidence intervals (HC3 robust standard errors). The dashed horizontal line indicates no difference (Δ = 0). Fiber types are displayed along the x-axis. Significant effects before multiple-testing correction are annotated with ‡. Effects that remain significant after multiple-testing correction (q-value) are shown: q<0.05: ★; q<0.01: ★★; q<0.001: ★★★; q<0.0001: ★★★★.

We found that ankle and elbow flexor-extensor muscle pairs had immature fiber type profiles in the sham rabbit P14 age group. Ankle extensors (gastrocnemius lateral head, gastrocnemius medial head, and plantaris) and elbow extensors (triceps brachii lateral head and triceps brachii long head) were highly fast-twitch oxidative. At this postnatal age, only a small subset of myofibers in these muscles were wholly glycolytic (type IIx, hybrid IIx/IIb, or IIb). The muscle with the fastest, most glycolytic profile was the gastrocnemius lateral head. In the P14 age group, sham flexor muscles, namely the biceps brachii (elbow flexor) and tibialis anterior (ankle flexor) were also largely devoid of the slowest (type I) and fastest (type IIx and type IIb) fiber types. The fiber type composition of the biceps brachii was: 7.4 % type I, 46.7 % type IIa, 42.0 % type IIa/IIx, 2 % type IIx, 1.2 % type IIx/IIb, and 0.8 % type IIb. The fiber type composition of the tibialis anterior was: 4.4 % type I, 39.7 % type IIa, 44.7 % type IIa/IIx, 5.0 % type IIx, 4.0 % type IIx/IIb, and 2.2 % type IIb (Table 1). The fiber type distributions that we measured in sham ankle and elbow extensors are typical of an immature neuromuscular system. While MyHC-I and MyHC-IIa are expressed from birth alongside fading expression of embryonic and neonatal myosins, MyHC-IIx is first detected at the beginning of the second postnatal week in rabbits and MyHC-IIb is the last adult fast MyHC to be expressed at the beginning of the third postnatal week, although the precise timing varies by muscle (Mckoy et al. 1998).

Postural muscles from sham rabbits, specifically the soleus and triceps brachii medial head, showed a balanced expression of slow-twitch and fast-twitch fibers in the P14 age group. This fiber type composition is characteristic of immature anti-gravity muscles, as muscles with this function are mixed-type at birth but become purely type I upon fast-to-slow fiber type transitions that occur during the first postnatal month (Larson et al. 2019; Wigston and English 1992). A complete report of the muscle fiber type compositions measured in this study can be found in Table 1.

We tested whether HI injury in utero affected muscle fiber type compositions in the P14 age group. We found HI injury induced a shift in muscle fiber composition specifically in flexor, but not extensor or anti-gravity muscles. HI flexor muscles had slower, more oxidative fiber type profiles than that of age-matched sham rabbits during the third postnatal week. Compared to that of shams, the elbow flexor biceps brachii and ankle flexor tibialis anterior of HI rabbits contained a greater proportion of type IIa fibers (biceps: +12% [95%CI-1%-+24%]; tibialis anterior:+22% [95%CI +5%-+39%], although these changes did not survive multiple comparison correction, at the expense of the proportion of hybrid type IIa/IIx fibers (biceps:-15% [95%CI-21%--8%], FDR-corrected p-value q=0.000254; tibialis anterior:-22% [95%CI-37%--6%], q=0.0997). Notably, we observed few differences in muscle fiber type compositions between HI rabbits with and without hypertonia as assessed by Ashworth scores (see methods). Specifically, the proportion of hybrid type IIa/IIx fibers was larger in the biceps brachii (+8% [95%CI +4%-+12%], q=0.0115) and smaller in the tibialis anterior (-12% [95%CI-14%--9%], q=7.72e-19) of hypertonic versus non-hypertonic HI rabbits. We also noted that the proportion of type I fibers in the plantaris was slightly greater in hypertonic HI rabbits (+1% [95%CI +1%-+2%], q=0.0328). Together, these results show that prenatal HI injury alters muscle fiber type composition, suggestive of a maturational delay in glycolytic fiber development.

This muscle fiber type shift was present in the HI group independently of hypertonia, demonstrating that muscle development was dysregulated by prenatal HI injury even when this insult was modest. This outcome suggests that motor deficits induced by prenatal HI injury may not be limited to hypertonia. In fact, data from our rabbit colony suggest an intermediate presentation of non-hypertonic HI kits, with locomotor speeds, time mobile, and total distance traveled in an open field falling intermediate between typically developing sham rabbits and hypertonic HI kits (Figure S1). Therefore, despite having normal muscle tone detected by Ashworth scoring, HI kits classified as non-hypertonic showed evidence of motor dysfunction.

We next studied fiber type compositions of the same flexor-extensor muscle pairs in the P31 age group, when kits are weaned and the neuromuscular system is functionally mature. Since only marginal differences in fiber type composition are reported between P30 and young adulthood in rabbits (Lobley et al. 1977), we chose to evaluate fiber type distributions at P30-32 when kits were pre-pubertal. In sham rabbits, ankle and elbow extensor muscles that were highly fast-twitch oxidative in the P14 group developed a major fast-twitch glycolytic component by weaning. The muscle with the fastest profile was again the gastrocnemius lateral head (26 % type IIb). As expected, the soleus and triceps brachii medial head, both anti-gravity muscles, transitioned from a mixed slow-and fast-twitch profile to a predominantly type I fiber type composition by weaning (soleus: 96 % type I; triceps brachii medial head: 79 % type I). The fiber type composition of the biceps brachii in P31 age groupe rabbits was: 8.0 % type I, 30 % type IIa, 16.8 % type IIa/IIx, 20.8 % type IIx, 12.6 % type IIx/IIb, and 11.8 % type IIb. The fiber type composition of the tibialis anterior was: 4.1 % type I, 23.3 % type IIa, 26.0 % type IIa/IIx, 30.5 % type IIx, 7.9 % type IIx/IIb, and 8.2 % type IIb. While fiber type compositions of the extensor and anti-gravity muscles that we evaluated were comparable between sham and HI rabbits in the P31 age group, HI flexor muscles retained a slower, more oxidative profile at weaning age. Again, this profile was not specific to hypertonic HI rabbits. Flexor muscles from P31 age group rabbits that experienced HI injury in utero were composed of a higher abundance of type IIa and hybrid IIa/IIx fibers (biceps: +8% [95%CI-0%-+17%], q=0.375 and +8% [95%CI +5%-+11%], q=1.25e-07, respectively; TA: +17% [95%CI +6%-+27%], q=0.0403 and +9% [95%CI +1%-+17%], q=0.235, respectively) at the expense of pure type IIx fibers (biceps:-16% [95%CI-22%--10%], q=1.48e-05; TA:-22% [95%CI-36%--9%], q=0.0251). The fiber type differences observed in HI flexor muscles in the P31 age group are unlikely to reflect immaturity because type IIb fibers are the last to develop, and the proportions of type IIb fibers were not significantly different between sham and HI in the biceps brachii (sham, 11.8 %; HI, 12.0 %; q=0.993) or tibialis anterior (sham, 8.2 %; HI, 5.7 %; q=0.826). Instead, this fiber type shift may reflect chronic, low-frequency motor unit activity.

### Muscle fiber area is unaffected by HI but increases with postnatal age

Since contractile force is positively correlated with myofiber size, we tested whether muscle fibers were smaller in young rabbits that experienced HI injury in utero. We evaluated muscle fiber cross-sectional area using a mixed-effects linear model (Figure 3A). A significant main effect of age revealed that for all rabbits, muscle fibers increased substantially in size (+131 % [95%CI +78%-+201%], p=4.79e-10) between P14 and P31 groups. Significant main effects of muscles (p<0.001) and fiber type (p<0.001) highlighted that average myofiber area is different across muscles and fiber physiological types. Myofiber cross-sectional area was smallest in the plantaris muscle and largest in the medial head of the triceps brachii. Type IIa fibers had the smallest cross-sectional area, and IIx/IIb hybrids were the largest (Figure S2).

**Figure 3:**
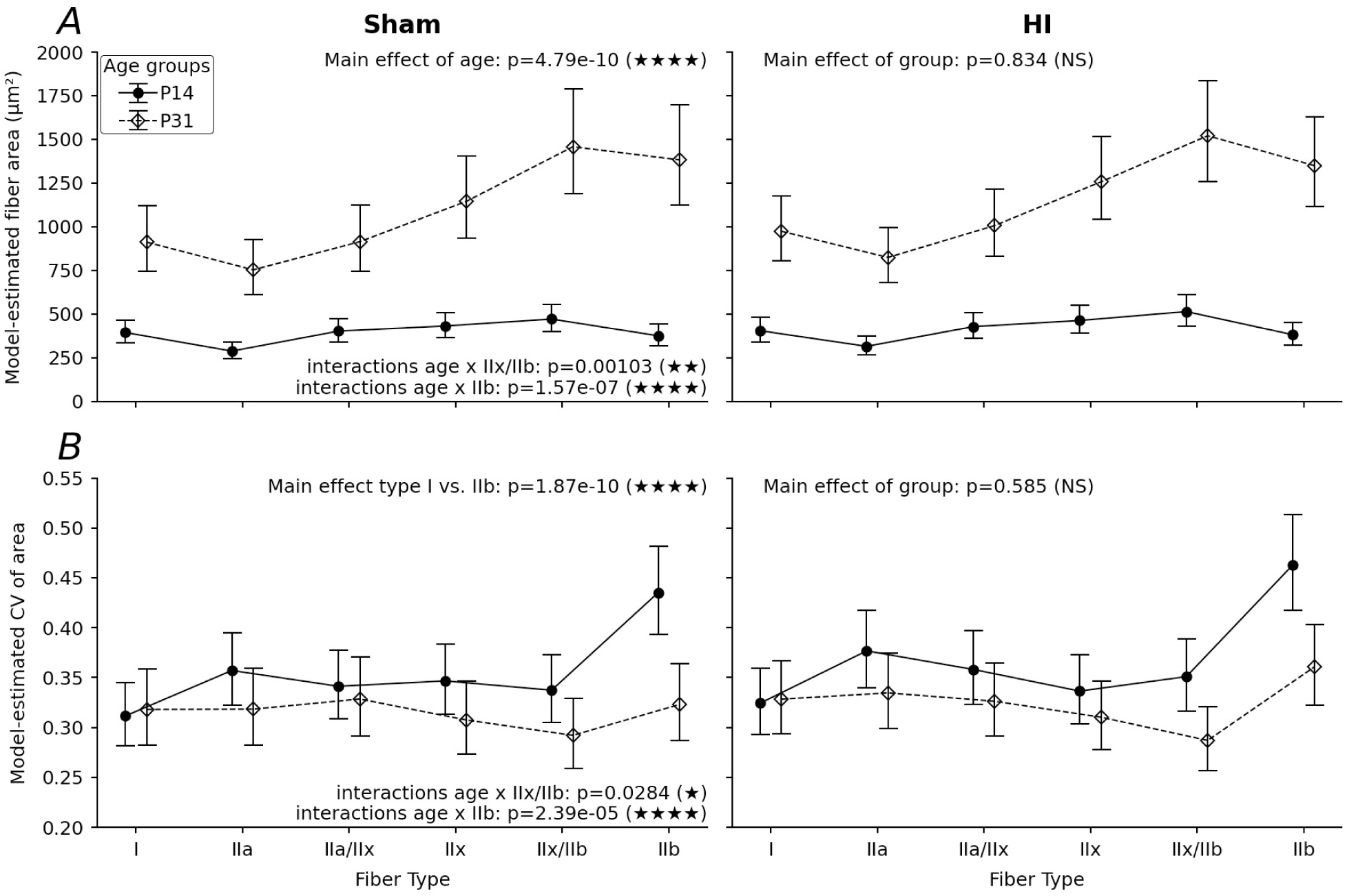
Fiber cross-sectional area (CSA) and size variability across age and treatment groups. **(A)** Estimated mean fiber cross-sectional area as a function of fiber type in Sham and HI animals at both time points. Points represent model-based estimates from linear mixed-effects models, and error bars indicate 95% confidence intervals. Solid lines and dots denote P14 animals and dashed lines and empty diamonds denote P31 animals. Fiber types are displayed along the x-axis. **(B)** Estimated coefficients of variability of fiber area as a function of fiber type, age and group. Same organization as in A. p values are extracted from our mixed-effect linear models. p<0.05: ★; p<0.01: ★★; p<0.001: ★★★; p<0.0001: ★★★★.

In our dataset, type IIa fibers were 27 % ([95%CI-35%--18%], p=3.35e-07) smaller than type I fibers; type IIx/IIb fibers were 19 % ([95%CI +6%-+35%], p=0.004) larger than type I fibers, while the area of other fibers were not significantly different than type I. The small size of type II fibers relative to type I fibers is a known hallmark of muscular immaturity in the rabbit (Gordon 1984), and is present in young humans as well (Esbjörnsson et al. 2021). Yet, there were significant interactions between age × fiber types, indicating that the sizes of hybrid IIx/IIb and pure type IIb fibers were disproportionately impacted by age; these fiber types hypertrophied much more between the P14 and P31 age groups compared to type I fibers (+34 % vs. growth in type I [95%CI +12%-+59%], p=0.00103 and +59 % vs. growth in type I [95%CI +34%-+90%], p=1.57e-07, respectively). The greater growth of fast-twitch glycolytic fibers (relative to type I fibers) between the P14 and P31 age groups reflects neuromuscular maturation. We did not observe a significant main effect of group (p=0.834), indicating that prenatal HI injury did not alter average myofiber size in the P14 age group. It is important to note, however, that muscle fiber cross-sectional area is directly related to body weight, regardless of muscle fiber type, (Esbjörnsson et al. 2021) and that rabbit body weight is heavily influenced by litter size (Poigner J. et al. 2010), which was uncontrolled in our study.

Variability of size in each type of myofiber is a morphometric property of muscle that is independent of weight; variation in myofiber CSA is unchanged between children and adults in non-disease contexts (Esbjörnsson et al. 2021). Increased fiber-size variation is indicative of myopathy (Valentine 2017) and may be positively correlated with functional motor deficits in CP (Edman et al. 2026; Pontén et al. 2005; Rose et al. 1994). Since it is unconfounded by variable body weight, we analyzed the co-efficient of variation of fiber size using a mixed-effects linear model (Figure 3B). We found a significant main effect of fiber type (p<0.001); directional analysis revealed that size variation was highest in fast fibers, with type IIb fibers being 40 % ([95%CI +26%-+55%], p=1.87e-10) more variable than type I fibers. We observed a significant age × fiber type interaction, showing that the most glycolytic fibers experience a larger decrease in variability with age than type I fibers (IIx/IIb:-15 % [95%CI-27%--2%], p=0.0284; IIb:-27 % [95%CI-37%--16%], p=2.39e-05) (Figure 3B and Figure S2). We did not detect a significant group × fiber type interaction (all p > 0.05), which demonstrated that fiber size variation was not increased in HI rabbits. Thus, we found that muscle fiber area and its co-efficient of variation were unaffected by prenatal HI injury but changed significantly with postnatal age.

## Discussion

In this study, we built a machine-learning model for automatic classification of muscle fiber type, and used this algorithm to quantify changes to muscle fiber type composition induced by prenatal HI injury, a risk factor for CP. Our XGBoost algorithm-based prediction model predicted muscle fiber type primarily by MyHC isoform immunofluorescence intensities, with minor influence from morphological descriptors (like roundness). Our model predicted pure fibers with high accuracy and handled hybrid fibers well. This fiber type classification model detected oxidative fiber type transitions in flexor muscles of young rabbits that experienced HI injury in utero (with and without hypertonia).

One major strength of our fiber type classification algorithm was its ability to measure hybrid fibers. Hybrid fibers express two MyHC isoforms, usually nearest neighbors along the physiological spectrum (Pette 2006; Pette and Staron 2001; Pette and Staron 1997). Hybrid expression can signify stable, intermediate contractile profiles or plasticity in the form of fiber type transitions, since MyHC protein turnover is slow (Medler 2019). In some muscles (but not others) hybrid fibers decline with postnatal age (Brummer et al. 2013; Larson et al. 2019). Hybrid fibers are a major obstacle to existing programs for semi-automatic classification of muscle fiber type. They are under-reported or not classified by existing programs, presumably because most hybrid fibers express MyHC-IIx plus another isoform (e.g., type IIa/IIx and type IIx/IIb fibers), and MyHC-IIx expression is defined in these programs as the absence of any fluorescent signal. MyHC-IIx expression is almost always inferred this way because of the limitations of multiplex microscopy; with only four imaging channels, the four MyHC isoforms plus one extracellular matrix marker cannot all be immunolabeled. The programs MuscleJ2 and QuantiMus circumvent this issue by immunolabeling for MyHC-IIx instead of MyHC-IIb, which works in limited circumstances, like when applied to rodent soleus or human muscle, which are devoid of type IIb fibers (Danckaert et al. 2023; Kastenschmidt et al. 2019). Hybrid fibers are very common; greater than or equal to one-fifth of murine, rabbit, and human muscle composition is estimated to be hybrid fibers (Medler 2019). Yet, hybrid fibers (or those with MyHC-IIx) are not considered by the MATLAB application SMASH (Smith and Barton 2014), the ImageJ macro from (Bergmeister et al. 2016), MyoVision (Wen et al. 2018), the script developed by (Reyes-Fernandez et al. 2019), the machine-learning algorithm Myosoft (Encarnacion-Rivera et al. 2020), MyoSight (Babcock et al. 2020), the Cellpose-Labels to ROIs pipeline (Waisman et al. 2021), MyoView (Rahmati and Rashno 2021), MuscleBos (Rehman et al. 2025), or FLASH (Di Gallo et al. 2025). The recently developed imaging analysis algorithm named MyoQuant (Madsen et al. 2025) reports “Hybrid” and “Hybrid/Other” fibers in the distribution but does not define their physiological types. Thus, we are the first to build a machine-learning algorithm for automatic classification of muscle fiber type that is able to identify common hybrids.

Our classifier measured fiber type compositions of ankle and elbow flexor-extensor muscle pairs at two unique stages in early postnatal development. In the P14 age group, all phasic muscles from sham and HI rabbits were composed of mostly fast-twitch oxidative fibers. Nonetheless, the fiber type compositions of flexor muscles (tibialis anterior and biceps brachii) were oxidative to an even greater degree in HI rabbits. They were composed of relatively more type IIa fibers and relatively fewer hybrid IIa/IIx fibers. Anti-gravity muscles (soleus and triceps brachii medial head) from both sham and HI rabbits expressed nearly equal proportions of slow-and fast-twitch fibers. At this developmental stage, muscle fiber size related to physiological type, but all fibers were much smaller in area than their counterparts at weaning age. These features highlight immaturity of rabbit musculature in the P14 age group and suggest that prenatal HI injury delays flexor muscle development.

Alternatively, muscle fiber type compositions and fiber morphology showed signs of maturity in flexor-extensor muscle pairs in the P31 age group. Phasic muscles contained a substantial fraction of entirely glycolytic fibers. This observation was expected, since glycolytic activity is known to increase over the first eight postnatal weeks in rabbits (Briand et al. 1993). Postural muscles had undergone fast-to-slow fiber type transition, a developmental hallmark of the first postnatal month (Larson et al. 2019; Wigston and English 1992). We observed proportions of type I and type II fibers in the tibialis anterior (4.1% type I; 95.9% type II) and plantaris (10.9% type I; 89.1% type II) that matched an early report inferred from Ca^2+^-ATPase activity in P30 rabbit muscle (Lobley et al. 1977): tibialis anterior: 5.3% type I; 94.7% type II; plantaris: 11.8% type I; 88.2% type II). However we report a greater proportion of type I fibers in the soleus (95.9% type I; 4.1% type II) compared to (Lobley et al. 1977)(86.9% type I; 13.1% type II), but our sampling approaches were different.

Here, we determined that flexor muscles remained more oxidative in HI rabbits at P30-32. Interestingly, this altered fiber type composition was observed in HI-injured rabbits with and without hypertonia based on Ashworth scoring. We surmise this was due to two main factors: (1) rabbits that displayed hypertonia in any joint as neonates were included in this category, but did not necessarily have hypertonia in all the muscle groups studied here, so some of the muscles from hypertonic rabbits were not themselves overtly hypertonic; (2) while severe cases of motor deficits and hypertonia are obvious, more subtle cases may be missed (particularly by subjective testing like Ashworth scoring), so some HI rabbits labeled as non-hypertonic may have had some degree of subtle hypertonia and/or other motor deficits (Figure S1). Here, in our HI rabbits, the tibialis anterior and biceps brachii contained higher proportions of type IIa and hybrid IIa/IIx fibers in lieu of type IIx fibers. The more oxidative (less fatigable) fiber type composition of flexor muscles in HI rabbits is likely permissive to sustained muscle activity and hypertonia.

Altered fiber type identity in the P31 age group could be reflective of delayed development, because oxidative fibers predominate at young ages. However, proportions of type IIb fibers were equal in sham and HI muscles despite the fact that MyHC-IIb is the last isoform to develop. Thus, delayed development seems unlikely. Instead, we suspect that this fiber type shift reflects a more universal, compensatory change adapting to more sustained motor unit activity in HI rabbits, likely associated with hypertonia. Motoneuron firing patterns influence muscle fiber type (Lømo et al. 1974; Salmons and Sréter 1976; Salmons and Vrbová 1969), and it has been demonstrated that chronic low-frequency stimulation causes fast-to-slow fiber type transitions in rabbits (Schuler and Pette 1996; Somasekhar et al. 1996). With this in mind, it is possible that the slower fiber type composition of flexor muscles in HI rabbits reflects chronic low-frequency motoneuron activity in these motor pools. This idea is supported by our prior studies linking prenatal HI injury to greater motoneuron excitability. We showed using in vitro spinal cord slices that motoneurons from hypertonic HI kits had greater sustained firing in artificial cerebrospinal fluid (Steele et al. 2020). HI likely increases motoneuron excitability by enhanced serotonergic neuromodulation, as spinal serotonin (5-HT) is elevated threefold in hypertonic HI rabbits (Drobyshevsky et al. 2015), and 5-HT is known to enable sustained motoneuron firing via 5-HT_2_ receptors (Harvey et al. 2006a; Harvey et al. 2006b; Ladewig et al. 2004). Indeed, HI motoneurons are more sensitive to neuromodulation by 5-HT, as they respond to 5-HT_1-2_ receptor agonism with a robust increase in excitability while control motoneurons do not (Reedich et al. 2023). Taken together, we postulate that elevated 5-HT levels in the spinal cord of HI rabbits may cause chronic motor unit activation in vivo.

Changes to the oxidative capacity of myofibers in affected muscles of people with CP have been reported. Both less glycolytic (Ito et al. 1996; Marbini et al. 2002) and more glycolytic (Deschrevel et al. 2024; Gantelius et al. 2012; Pontén et al. 2008) profiles have been documented. One possible explanation for this inconsistency is that previous treatments like immobilization or Botox injections may have impacted the results. In fact, both immobilization and Botox are known to alter myofibers to cause slow-to-fast fiber type transitions (Valentine et al. 2016). Interestingly, in a well-controlled study of the erector spinae, which is a postural muscle as opposed to other studies which targeted limb muscles that were impacted by the condition, a fast-to-slow fiber type transition was observed as well (Robinson et al. 2013). Our current findings are consistent with prior studies reporting glycolytic-to-oxidative fiber type transitions; however, we note fiber type transitions are highly muscle-specific and there are few clinical reports studying the muscles we examined here. A shift in the proportion of fatigable muscle fibers may contribute to weakness in people with CP, as fatigable muscle fibers provide more power during contractions. It could also impact orderly recruitment during contractions, as the oxidative (fatigue-resistant) fibers are recruited before the fatigable glycolytic fibers. In sum, altered muscle fiber type compositions could contribute to motor deficits in CP.

Increased variability in muscle fiber CSA is observed in muscle biopsies from individuals with CP (Deschrevel et al. 2024; Deschrevel et al. 2026; Kahn et al. 2023; Rose et al. 1994), but we did not observe this phenomenon in HI rabbits in either the P14 or P31 age groups. Inconsistencies between studies are problematic, and make interpretation difficult. In human studies, it is more difficult to control variables including previous treatments like immobilization or Botox (which is known to alter myofibers to more glycolytic types), which muscles are included in the study (only those that are affected by the condition, or a consistent muscle / muscle group), how the muscles are impacted (spasticity vs. dystonia, constant vs. intermittent activity), and demographics like age, sex, weight, and gross motor function classification system (GMFCS) level plus availability of the participants. While we are able to conduct the present animal study using in many ways a more controlled approach, we also faced the challenge of variability in the muscles impacted by prenatal HI injury, and variations in body weight / litter size of the rabbits. Another limitation of our study is rooted in species-specific differences in muscle physiology, which are important to consider when comparing fiber type compositions of human versus rabbit muscle. Human muscle does not express MyHC-IIb and thus has a significantly slower profile than that of the rabbit or mouse, with rabbits being more similar to humans than mice (Bottinelli and Reggiani 2000; Ennion et al. 1995; Smerdu et al. 1994).

## Conclusions

In summary, this study developed a novel algorithm for high-throughput classification of muscle fiber type, including hybrid fibers, which could be used in many other studies. We provide evidence that sustained motor unit activity after prenatal HI injury drives a glycolytic-to-oxidative shift in flexor muscle fiber type composition at P14-20 and P30-32. This altered fiber type profile likely imparts less fatigability and reduced force-generative capacity to flexor muscles, which could contribute to motor dysfunction after prenatal HI injury.

## List of abbreviations

Botox: Botulinum neurotoxin type-A
BPM: Breaths per minute
CP: Cerebral palsy
CSA: Cross-sectional area
E: Embryonic day
GMFCS: Gross Motor Function Classification System
HI: Hypoxia-ischemia
Lat. Gast.: Gastrocnemius lateralis muscle
Med. Gast.: Gastrocnemius medialis muscle
MyHC: Myosin heavy chain
*MYH1*: *Myosin heavy chain 1*
*MYH2*: *Myosin heavy chain 2*
*MYH4*: *Myosin heavy chain 4*
*MYH7*: *Myosin heavy chain 7*
P: Postnatal day
PBS: Phosphate-buffered saline
PBST: Phosphate-buffered saline with 0.1% Tween-20
Plant.: Plantaris muscle
SHAP: Shapley Additive Explanations
Tib. Ant.: Tibialis anterior muscle
Triceps Brac. Lat.: Lateral head of the triceps brachii muscle
Triceps Brac. Long: Long head of the triceps brachii muscle
Triceps Brac. Med: Medial head of the triceps brachii muscle
WGA: Wheat germ agglutinin
5-HT: Serotonin

## Declarations

### Ethics approval

All experimental procedures involving animals were approved under protocol AN1718-001 by the University of Rhode Island’s Institutional Animal Care and Use Committee and were carried out in accordance with the National Institutes of Health Guide for the Care and Use of Laboratory Animals.

### Consent for publication

Not applicable

Availability of data and materialsThe data and analysis pipeline presented in this study are openly available in Zenodo at: https://doi.org/10.5281/zenodo.20602640.

### Authors’ contributions

CK: investigation, methodology, validation, visualization, writing - original draft, writing - review & editing. ER: conceptualization, investigation, methodology, validation, visualization, writing - original draft, writing - review & editing. HM: investigation, methodology, validation, visualization. SD: investigation. DS: conceptualization, investigation, methodology, visualization. EG: investigation. TU: investigation, visualization. AM: investigation, visualization. EMA: investigation. BM: investigation. LG: investigation, visualization. JG: investigation, visualization. CQ: investigation. LD: investigation. CJD: methodology. KQ: conceptualization, funding acquisition, investigation, methodology, writing - original draft, writing - review & editing. MM: conceptualization, data curation, formal analysis, funding acquisition, investigation, methodology, project administration, software, supervision, validation, visualization, writing - original draft, writing - review & editing.

## Acknowledgements

We gratefully acknowledge Amanda Dowst and the rest of the staff and resources of the University of Rhode Island Comparative Biology Resources Center for their expert animal care, technical assistance, and research environment that made this work possible.

Research was made possible by the use of equipment available through the Rhode Island Institutional Development Award (IDeA) Network of Biomedical Research Excellence from the National Institute of General Medical Sciences of the National Institutes of Health under grant number P20GM103430 through the Centralized Research Core facility and/or the Molecular Informatics Core (RRID:SCR_017685). Cryostat used to prepare some of the cryosections in this work is supported by the National Institute of Health equipment grant S10 OD032209.

The computational resources for this work were provided by the University of Rhode Island’s partnership with the Unity Research Computing Platform, a multi-institutional cluster led by the University of Massachusetts and the University of Rhode Island.

## Authors’ information

The contribution of CJD to the current study’s conceptualization was made at her previous institution (Lurie Children’s Hospital of Chicago and Northwestern University), not at NIH. The opinions expressed in this article are the author’s own and do not reflect the view of the National Institutes of Health, the Department of Health and Human Services, or the United States government.

---

**Supplemental Figure S1:**
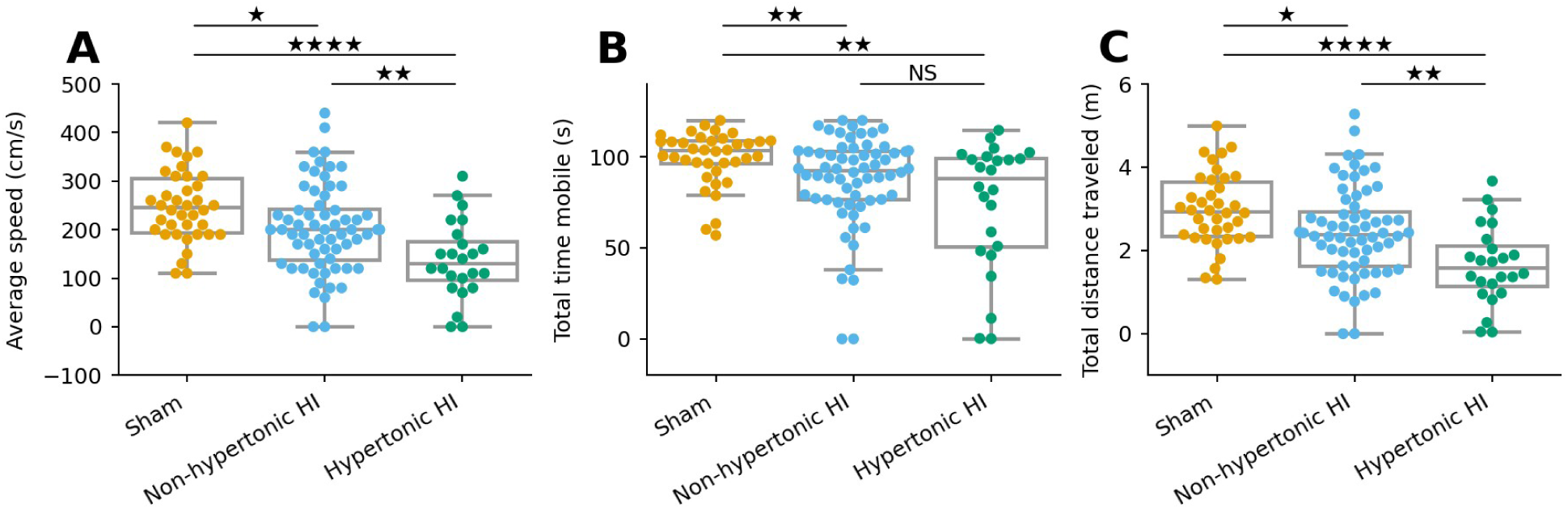
Measures of general locomotor ability of neonatal rabbits in an open field at P1. **(A)** Average speed, **(B)** total time mobile, and **(C)** total distance traveled of neonatal rabbits for the duration of the open field assay. Points represent individual rabbits (shams are orange circles, non-hypertonic HI are blue circles, hypertonic HI are green circles), superimposed over a boxplot that shows the three quartile values of the distribution. The “whiskers” extend to points that lie within 1.5 x inter-quartile distance of the lower and upper quartile. p<0.05: ★; p<0.01: ★★; p<0.0001: ★★★★

**Supplemental Figure S2:**
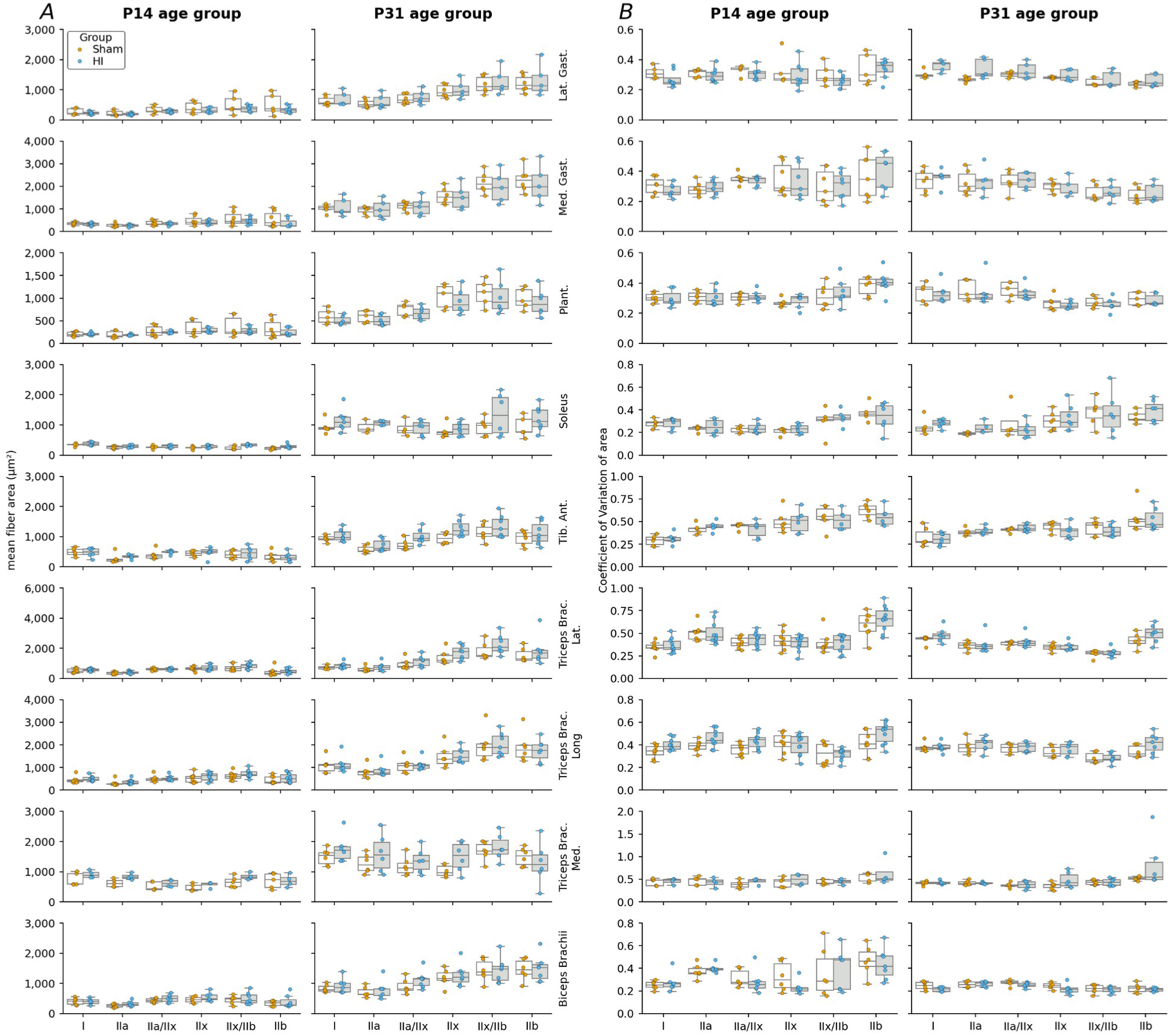
Fiber cross-sectional area and size variability across age, treatment groups and muscles. **(A)** Average cross-sectional area as a function of fiber type (x-axis), experimental group (Sham: left orange points, HI: right blue points), age (columns) and muscles (rows). Each point represents an animal. The boxplots show the three quartile values of the distribution along with extreme values. The “whiskers” extend to points that lie within 1.5 x inter-quartile distance of the lower and upper quartile. **(B)** Same representation for the coefficient of variation of the fiber areas.

