## Supplementary figures and images for "A novel machine-learning classification model detects oxidative fiber type transitions in a rabbit model of cerebral palsy"

### Figure S1

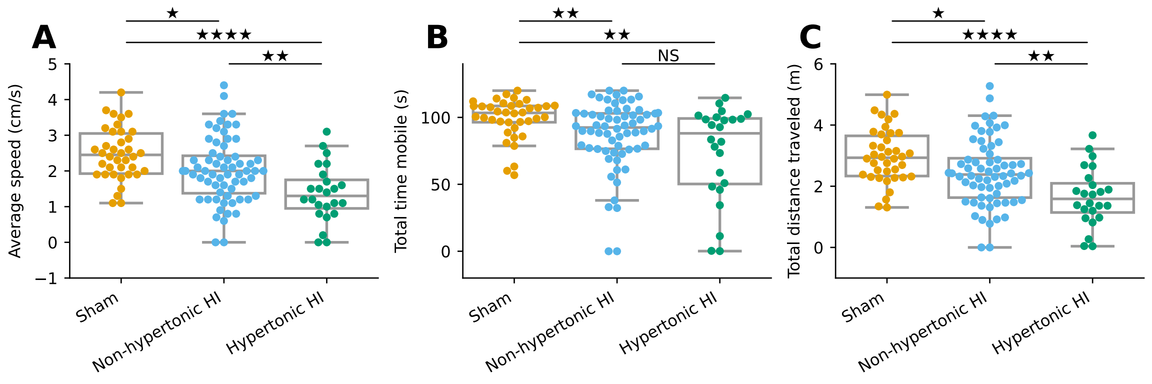

### Figure S2

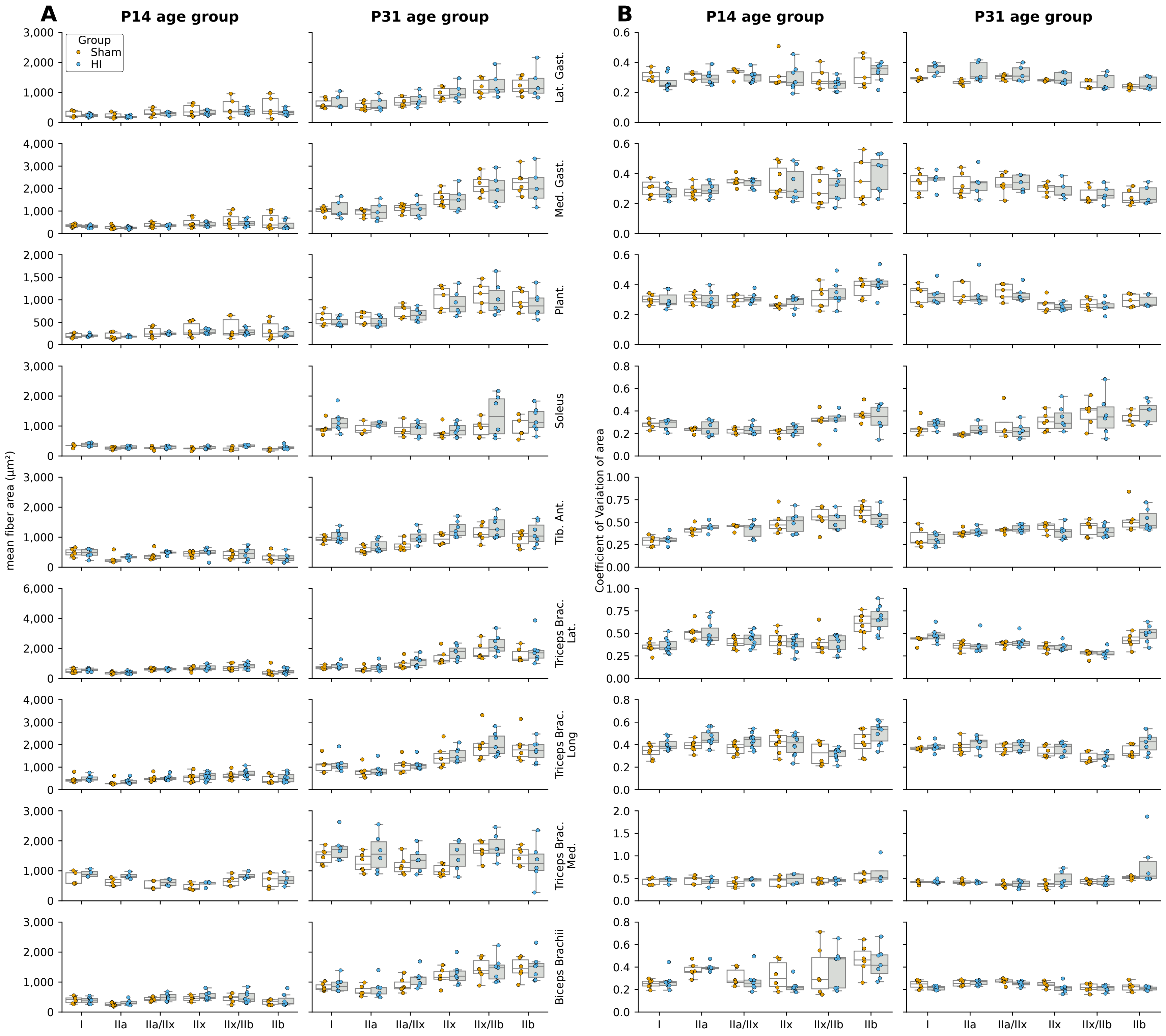
